# Biophysical properties of a tau seed

**DOI:** 10.1101/2021.03.30.437772

**Authors:** Zhiqiang Hou, Dailu Chen, Bryan D. Ryder, Lukasz A. Joachimiak

## Abstract

Pathogenesis of tauopathies involves conversion of tau monomer into pathological tau conformers that serve as templates to recruit native tau into growing assemblies. Small soluble tau seeds have been proposed to drive pathological tau assembly *in vitro*, in cells and *in vivo*. We have previously described the isolation of monomeric pathogenic tau seeds derived from recombinant samples and tauopathy tissues but in-depth biophysical characterization of these species has not been done. Here we describe a chromatographic method to isolate recombinant soluble tau seeds derived from heparin treatment. We used biochemical and biophysical approaches to show that the seeds are predominantly monomeric and have the capacity to nucleate aggregation of inert forms of tau *in vitro* and in cells. Finally, we used crosslinking mass spectrometry to identify the topological changes in tau as it converts from an inert state to a pathogenic seed. Future studies will reveal the relationship between soluble seeds and structural polymorphs derived from tauopathies to help diagnose and develop therapeutics targeting specific tauopathies.

## Introduction

Under normal physiologic conditions, tau is stable and does not readily aggregate in the absence of inducers [1]. Early analyses of tau structure suggested that it does not adopt a folded conformation, but rather is intrinsically disordered [2]. Given that tau encodes amyloid motifs that mediate self-assembly [3], a key question is what limits the self-association of amyloid motifs to yield aggregation-resistant tau under normal physiologic conditions? Recent isolation and characterization of distinct pools of tau monomer, some with properties of seeding and self-assembly, and others without, indicates that tau adopts a structure surrounding the amyloid motifs that regulates aggregation [4].

Prior studies have proposed that repeat domains encode local structure [5]. Tau has been shown to adopt structure surrounding the PGGG motifs, at the end of each repeat, that precede amyloid motifs [6]. Disease-associated mutations are enriched upstream of the amyloid motifs [7]. We have characterized the sequences surrounding the ^306^VQIVYK^311^ amyloid motif, finding that, disease-associated mutations upstream from this motif promote aggregation of tau by disrupting the protective local structures [3]. Thus, the formation of these protective structures limits aggregation but is still compatible with conformations required for association with microtubules. A model based on local structures that mask aggregation properties could explain tau’s stability and inability to aggregate in the absence of inducers. Interestingly, similar concepts have emerged for other intrinsically disordered proteins encoding amyloid motifs such as α-synuclein, where design of different β-turn types modulates aggregation properties [8].

Initiation of tau aggregation *in vitro* requires the addition of preformed tau seeds or incubation with polyanions, such as heparin, disrupt these aggregation-protective structures [9, 10]. Heparin binds to the repeat 2 in the repeat domain of tau, stabilizing it in an unfolded conformation [11-13]. Polyanion binding to positively charged residues in the repeat domain may preferentially expose sequences that promote oligomer assembly during the lag phase followed by the elongation adhering to a classical nucleation mechanism [14]. In prior work, we isolated distinct pools of tau monomer, from both recombinant and brain-derived sources, indicating that tau may exist in distinct separable conformations with different aggregation properties. Tau monomer that is otherwise aggregation-resistant has the capacity to adopt aggregation-prone conformations that self-assemble and initiate aggregation *in vitro* and in the setting of disease states [4]

Structural and modeling analyses comparing inert and seed-competent tau monomers revealed preferential exposure of amyloid motifs (^275^VQIINK^280^ and ^306^VQIVYK^311^) in seed-competent tau monomer which then can act as a nucleus to promote elongation [4]. Furthermore, the seed-competent form of tau monomer isolated from distinct tauopathies has been observed to encode distinct subsets of strains. This indicates a possible ensemble of aggregation-prone monomeric conformers that have the capacity to adopt and propagate distinct fibrillar conformations [15]. The idea that tau monomer alone can drive its assembly and serve as a template to form structural polymorphs is not widely accepted, although recent work from other groups on tau [16], and Sup35 [17], are consistent with this idea.

Here we describe a chromatographic approach to produce monomeric heparin-induced tau seeds, herein referred to as M_s_. We used cell-biological and biochemical approaches to study the properties of the M_s_ *in vitro* and in cells revealing that nanomolar amounts of seeds can trigger aggregation of inert tau (herein referred to as M_i_). We also show using biophysical approaches that M_i_ is a stable monomer at low or high concentrations while M_s_ is predominantly a monomer at nanomolar concentrations but at micromolar concentrations, it exists as an equilibrium between a monomer and dimer. Finally, we used a structural approach to highlight changes in topology along the pathway to tau seed formation revealing structural rearrangements that involve the acidic N-terminus through the N1/N2 domains, the basic proline-rich domain (P1 and P2), and the repeat domain (RD). Our data support that brief incubation of tau with heparin following chromatographic separation produces a seed that is predominantly a monomer, which has the capacity to nucleate tau aggregation *in vitro* and in cells; and the monomeric seed displays distinct conformations. A deeper understanding of the conformation of tau seeds will yield insight into the mechanisms of structural polymorph formation and how ligands or cofactors can mediate this conversion into pathogenic forms of tau.

## Results and Discussion

### Heparin-based conversion of tau into a pathogenic seed

Initial efforts to produce small soluble tau seeds, including monomer, were based on sonication of tau fibrils followed by size exclusion chromatography (SEC) to separate the different sized species [4, 18]. *In vitro* and in cell seeding assays confirmed the aggregation behavior of these small soluble species. Additionally, similarly sized species from different tauopathies were isolated from brain tissues [19, 20]. Here we developed a method to consistently produce recombinant monomeric tau seeds without fibril sonication. Typically, tau aggregation involves incubation of tau in the presence of polyanions such as heparin in buffered saline. We discovered that incubation of these reactions in a sulfated buffer, such as 3-(N-morpholino) propanesulfonic acid (MOPS), slows the conversion of tau into larger oligomers allowing us to resolve small soluble species by SEC before they convert into large oligomers (Fig 1a). We observed that on a Superdex 200 SEC column (GE), normal inert tau (herein M_i_) elutes as a monodisperse peak (Fig b; blue trace, 13.5mls) while tau incubated with heparin elutes earlier (Fig 1b; red trace, 12mls). To first evaluate the capacity of these tau species in mediating aggregation, we employed HEK293T tau biosensors that express tau repeat domain fused to CFP and YFP. This in-cell aggregation assay is sensitive, specific, and can detect tau aggregates down to femtomolar concentrations [21]. Transduction of negative (lipofectamine alone) and positive controls (recombinant tau fibrils) yielded expected signal with 0.14% ± 0.10 and 81.47% ± 4.32, respectively (Fig 1c). Lipofectamine transduction of 100nM M_i_ reisolated from SEC (i.e. Fig 1b, blue trace, 13.5ml fraction) into biosensors cells yielded 0.17% ± 0.11 cells with aggregates (Fig 1c), similar to lipofectamine alone. Transduction of 100nM tau reisolated from the tau:heparin reaction (i.e. Fig 1b, red trace, 12ml fraction) yielded 39.47% ± 0.55 of cells with aggregates (Fig 1c). Given that the “seed” eluted earlier on an SEC column, we wondered whether this tau species is still bound to heparin. To monitor the heparin on the SEC, we utilized a heparin-fluorescein (FITC) conjugate allowing facile tracking of the absorbance signal at 488nm. Heparin-FITC alone eluted at 20.5mls (Fig 1b; yellow trace). The heparin-FITC absorbance for the tau:heparin-FITC reaction eluted as an overlapping peak with heparin-FITC alone (Fig 1b; purple trace) with no detectible 488nm signal in the fraction which induced tau aggregation in cells. To independently confirm the concentration of heparin-FITC signal in the M_s_ peak fraction (Fig 1b, purple, 13mls), we produced a fluorescence calibration curve for FITC. We can detect FITC fluorescence signal down to 10e^-13^ M and it appears linear in the range from 0.05 nM to 1 μM (Supplementary Figure 1a). Measurement of FITC fluorescence in the 13.5ml peak from the tau:heparin reaction yielded an estimated 4 nM FITC signal while the A205 protein signal yields a tau concentration of 4 μM. This suggests that SEC is quite efficient at separating the heparin away from the tau leaving only trace amounts that are on the order of 1:1000 (Supplementary Figure 1a). Our data support that brief incubation of recombinant tau with heparin followed by re-isolation by SEC produces small molecular-weight tau seeds that contain negligible amounts of heparin.

**Figure 1.**
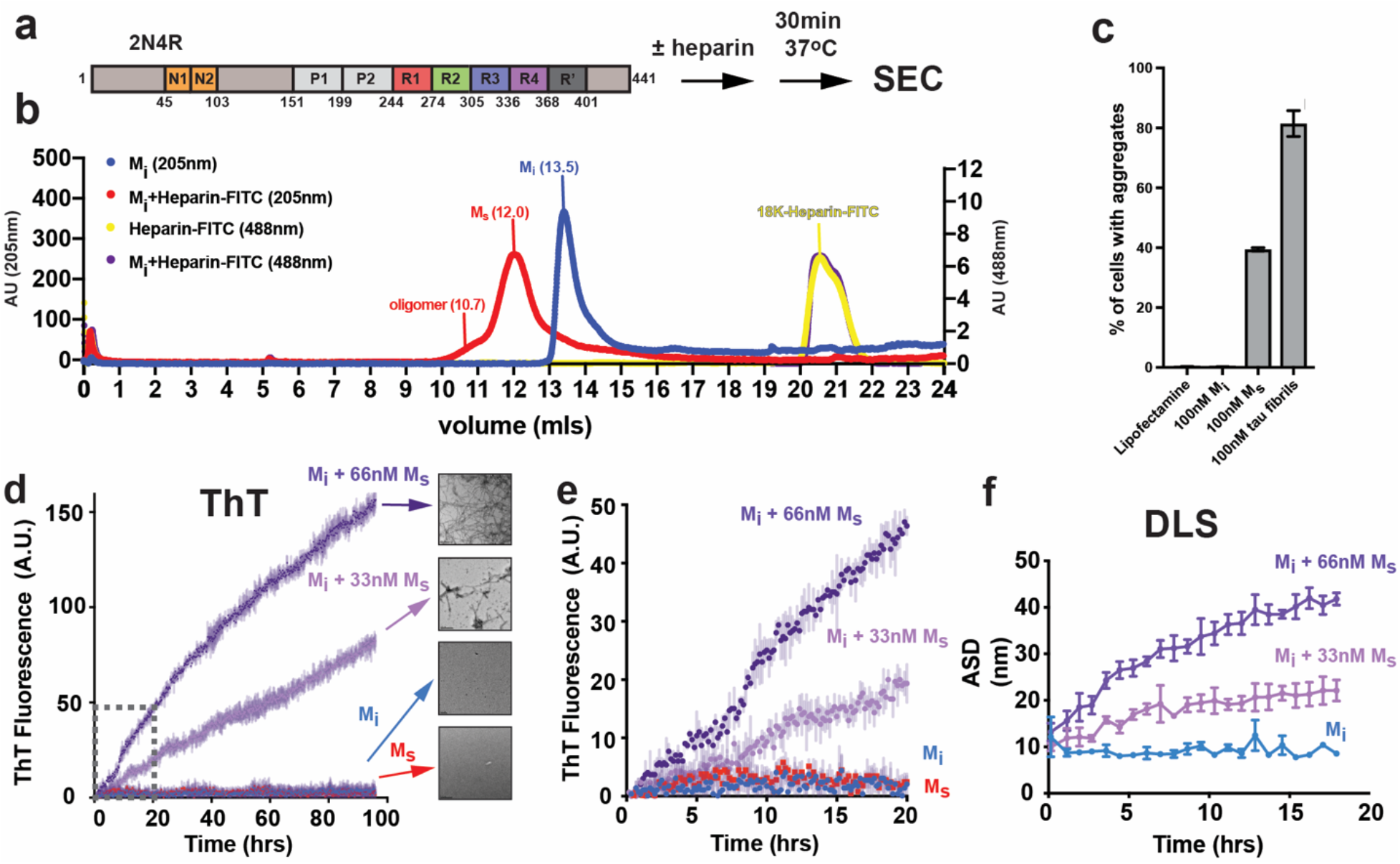
Generation and characterization of recombinant tau seeds. **a** Schematic for tau seed production. Full-length 2N4R tau was incubated with heparin for 30 minutes and resolved by SEC. 2N4R tau is shown as a cartoon schematic. The N1 and N2 domains are colored in orange. The repeat domains are colored in red, green, blue, magenta and dark grey. The proline-rich P1 and P2 domains are colored in light grey. **b** SEC chromatograph of tau, tau:heparin-FITC, heparin-FITC reactions resolved on a Superdex 200 10/300 GL increase column. The tau alone and tau:heparin-FITC traces acquired using A205 are colored in blue and red, respectively. The heparin-FITC and tau:heparin-FITC traces acquired using A488 are colored in yellow and purple, respectively. **c** Activity of SEC fractions in a tau biosensor seeding assay. Lipofectamine alone and tau fibrils were used as negative and positive controls, respectively. Experiments were performed in triplicate showing average values with standard deviation. **d** ThT fluorescence aggregation assay comparing 4uM M_i_ (blue), 66nM M_s_ alone (red), 4uM M_i_ + 33nM M_s_ (light magenta), 4uM M_i_ + 66nM M_s_ (purple). Representative TEM images of M_i_, M_s_, M_i_ + 33 nM M_s_ and M_i_ + 66 nM M_s_ samples imaged at the end point of each reaction. Fibrils were not observed in the M_i_ alone condition. Each experiment was performed in triplicate and is shown as an average with standard deviation. Grey dotted box highlights the data for the early 20 hour time point for comparison with DLS data. **e** Zoom in of ThT fluorescence aggregation experiment from Figure 1D within 20hrs allowing direct comparison of the fluorescence signal to the DLS experiment in **f**. Curves are colored as in Figure 1D. **f** DLS time-course of seeded M_i_ aggregation. Average size distribution (ASD) of triplicate 4uM M_i_ alone (blue), 4uM M_i_ + 33 nM M_s_ (light magenta) and 4uM M_i_ + 66n (purple). 66nM M_s_ alone was not included because it was to dilute to observe by scattering. DLS experiment was carried out in triplicate and the data are shown as averages with standard deviation.

We wanted to show that our recombinant tau seeds are capable of inducing M_i_ aggregation using an *in vitro* Thioflavin T (ThT) fluorescence aggregation assay. Addition of 33 nM or 66 nM tau seeds to 4 μM M_i_ tau yields a robust increase in ThT fluorescence while M_i_ alone and M_s_ alone controls remained flat. We confirmed the presence of tau fibrils at the end of the reactions using Transmission Electron Microscopy (TEM). Consistent with the ThT experiments, inert tau samples treated with 33 nM or 66 nM seeds yielded fibrils by TEM while M_i_ or M_s_ alone did not (Fig 1d,e). ThT fluorescence aggregation assays are the gold standard to detect β-sheet rich amyloid structures *in vitro* and we hypothesized that tau assembly can be monitored using dynamic light scattering (DLS). As in the ThT assay, M_i_ alone remained stable over the course of the 20hr experiment with an average size distribution (ASD) of 10nm (Fig 1f) skewed by a <0.1% fraction of large species. Indeed, binning the data across sizes revealed predominantly a distribution of sizes ranging from 1-7nm centered at 2.4nm with no significant fraction of larger species (Supplementary Figure 1b). Addition of 33 nM or 66 nM tau seeds to 4 μM M_i_ begins with an ASD around 10nm (also skewed by <0.1% larger species) and grows steadily over 20 hours to 20nm and 40nm, respectively (Fig 1f). Binning the sizes of these samples at t=0hr showed similarity to the M_i_ alone condition, each seeded experiment predominantly starts as a distribution of sizes from 1-8nm centered at 2.4nm (Supplementary Figure 1c,d, green). After 20 hrs in the 33 nM seeded samples, the smaller species shift to 4.2nm and larger oligomers centered on 41.2nm are now present (Supplementary Figure 1c, red). After 20hrs in the 66nM seeded samples, the sizes shift yet further to species centered on 5.6nm and 72.6nm (Supplementary Figure 1d, red). The DLS assay allows us to monitor the conversion of tau into larger species by following average sizes across the entire distribution but also to quantify the distribution of small and large oligomers over time. Interestingly, we can detect tau assembly using DLS at early time points where the ThT fluorescence is low and unreliable (Fig 1e). suggesting this is a good orthogonal assay to detect tau assembly early in aggregation reactions prior to robust ThT signal.

### Quantification of tau seed size

To confirm the size of the heparin-induced tau seed species, we employed biophysical approaches including DLS, Size Exclusion Chromatography Multi-Angle Light Scattering (SEC-MALS) and mass photometry (Fig 2a). First, we measured the size of M_i_ and the tau seed, M_s_, using DLS each at 4 μM. Our analysis suggests that the M_i_ size is small ranging in size between 1-7nm and centered on 2.4nm (Fig 2b, blue) while the M_s_ tau seed has a narrower size distribution from 3nm-5nm centered on 3.5nm suggesting a larger size for the seed (Fig 2b, red). While light scattering methods require high concentration samples, it is a method to unambiguously determine the molecular weight of protein species. Analysis of 44 μM M_i_ by SEC-MALS revealed good agreement with monomer with a molecular mass of 47.9kDa (Fig 2c). In contrast, analysis of 44 μM recombinant tau seeds induced by heparin revealed them to be a mixture of monomer (47.9kDa) and dimer (94kDa) with a minor population of oligomers eluting around the void volume (Fig 2d). Capturing the true size of these seeds given their capacity to drive assembly under much lower concentration is challenging given the necessary concentration requirements (∼2mg/ml) for SEC-MALS. To circumvent this issue, we employed a mass photometry approach that utilizes the principles of interference reflection and scattering microscopy to measure molecular weight of samples in solution at dilute concentrations. We used this method to determine the molecular mass of our tau species using 100nM concentrations, several orders of magnitude lower than required for SEC-MALS. We found that M_i_ is a monomer with a molecular weight of 49kDa fitting 97% (σ=7) of the population (Fig 2e). Analysis of the tau seeds showed that 83% of the population is 49kDa (σ=7) with 11% of the population as a dimer with a molecular weight of 91kDa (σ=9) (Fig 2f). These analyses suggest that the seeds are indeed aggregation-prone and at high concentrations equilibrate between monomer and dimer while at low concentrations they remain predominantly monomer.

**Figure 2.**
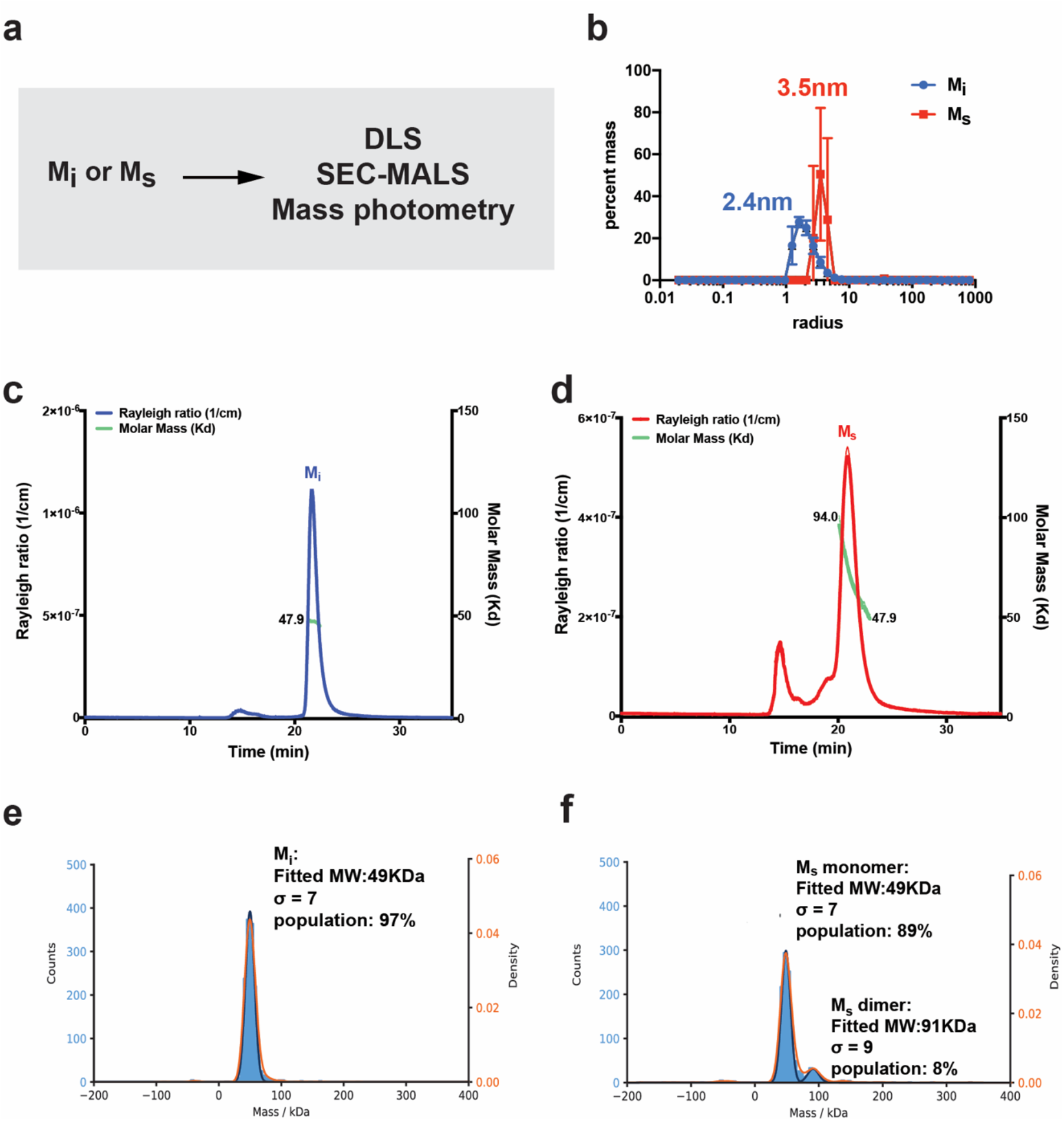
Quantification of tau seed shape and mass. **a** Schematic for experiments to measure the size of M_i_ and M_s_ using DLS, SEC-MALS and mass photometry. **b** Histogram of sizes observed for 4uM M_i_ (blue) and 4uM M_s_ (red) in DLS. **c** SEC-MALS of 16uM M_i_ (blue) shows a single peak that was calculated to have a molar mass of 47.9 g/mol (green). The single peak indicates that M_i_ tau elutes as a monomer. **d** SEC-MALS of 16uM M_s_ (red) shows a broader peak that was calculated to range from 94 g/mol to 47.9 g/mol (green). The broader peak indicates that M_s_ tau elutes as a distribution of dimer and monomer with minor signal from oligomers. **e** Mass photometry measurements of 50 nM M_i_ reveals the sample to be uniformly monomeric with a calculated molecular weight of 49kDa accounting for 97% of the sample with a sigma of 7. **f** Mass photometry measurements of 50 nM M_s_ reveals the sample to be predominantly monomeric with a calculated molecular weight of 49kDa accounting for 89% of the sample with a sigma of 7. We also observe a small proportion of a dimer with a calculated molecular weight of 91kDa accounting for 8% of the sample with a sigma of 9.

### Tau conformational changes in seed formation

Our prior studies on M_i_ and tau seeds involved crosslinking mass spectrometry (XL-MS) to understand possible changes in topology between these two conformations [3, 4]. Our first experiments on tau employed a homo bi-functional disuccinimide suberate (DSS) crosslinker which reports on contacts between lysine residues [22]. We have recently employed a new chemistry that can report on zero-length crosslinks directly between lysine residues and amino acids containing carboxylic acids mediated by 4-(4,6-Dimethoxy-1,3,5-triazin-2-yl)-4-methylmorpholinium chloride (DMTMM). In full-length tau we observed contacts between the acidic N-terminus and the basic repeat domain and we showed that pathogenic mutations including P301L alter the distribution of contacts from the N-term to the repeat domain [22]. Building on these ideas, we wondered how the N-term contacts to the repeat domain in full-length tau change from M_i_ to a seed. We reacted Mi, M_i_:heparin and M_s_ with DMTMM for 15 minutes, the samples were resolved by SDS-PAGE (Supplementary Figure 2a-c) and the monomer bands were extracted from the gel. The samples were processed using our XL-MS pipeline to identify changes in crosslink contacts with regard to XL sites and frequencies in these three different states. We observe 72 crosslinks in M_i_ (Fig 3a; blue) compared to 40 and 39 for M_i_:heparin and M_s_, respectively (Fig 3a; orange and purple). Additionally, we find that the M_i_ contacts are more heterogeneous, which are consistent with tau being an intrinsically disordered protein. Meanwhile, the M_i_:heparin and M_s_ contacts exhibit more reproducible modes of contacts (Supplementary Figure 2d-f). Consistent with our previous observations in full-length 2N4R tau [22], in M_i_ we observe 14 and 15 contacts from the N-terminal acidic N1/N2 domains to Proline-rich Domain 1 (P1) and Proline-rich Domain 2 (P2)/RD, respectively (Fig 3ab). In the M_i_:heparin complex, we observe 7 contacts from the N-term N1/N2 to P1 but a complete loss of contacts from the N-term N1/N2 to P2/RD (Fig 3ac). In M_s_, which is repurified from heparin incubation using SEC we recover 15 contacts from the N-term N1/N2 to P1 (Fig 3ad) but only 3 contacts from the N-term to P2/RD are recovered (Fig 3ad). Notably, we also observe a loss of crosslinks in the C-terminus in M_i_:heparin and M_s_ relative to M_i_ (Fig 3b-d). These data indicate that in the M_i_:heparin complex, the polyanion binds to the RD and displaces contacts from the acidic N-term N1/N2 to basic P1/P2 and RD. In M_s_ conformation, where the heparin has been removed, leaves the RD more exposed. In addition to crosslinks, our XL-MS data also yield information about solvent accessibility by monitoring the crosslinker reactions to a single amino acid which can be used as a measure of solvent accessibility. Consistent with the crosslinking data, we observe a higher incidence of adipic acid dihydrazide (ADH) monolinks (i.e. solvent exposure) at the acidic N-term N1/N2 in the M_i_:heparin complex compared to the M_i_ and M_s_ samples (Supplementary Figure 2g-i). Importantly, we have previously proposed that local protective structures engage amyloid motifs to prevent their self-assembly and consistent with this model heparin may play a role in disrupting both local structure and displacing long-range aggregation-protective contacts from the acidic N-term N1/N2 to P1 and P2/RD.

**Figure 3.**
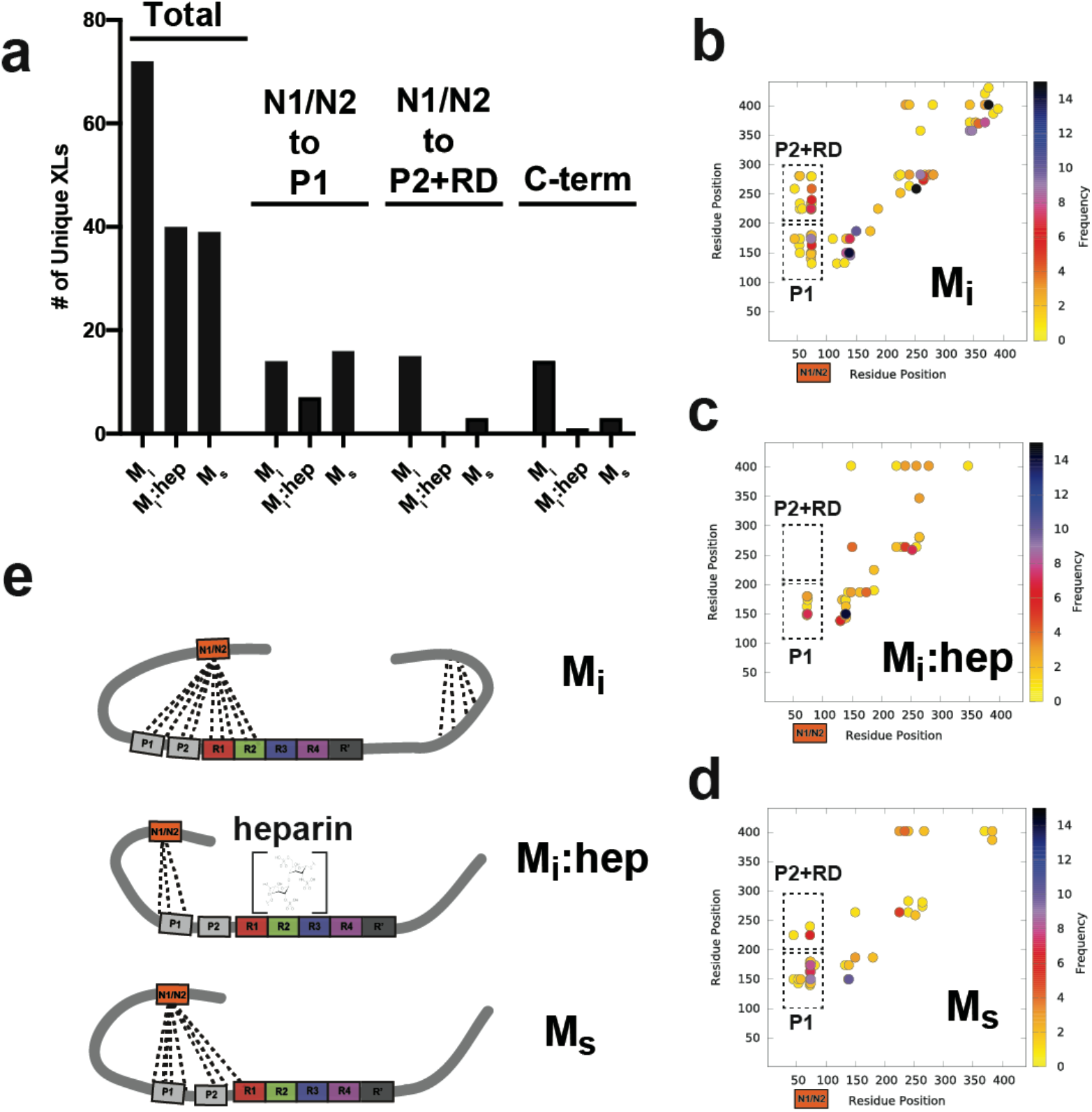
Topological changes in tau during seed formation. **a** Summary of unique crosslinks identified in M_i_, M_i_:heparin and M_s_. Experiments were performed as five replicates and data is shown as unique crosslinks observed across replicates. Total crosslinks are shown on the left, specific crosslinks observed from the acidic N-terminal N1/N2 domains (residues 45-101) to the P1 domain (residues 151-198, middle) and P2/RD domains (residues 199-368, right). Scatter plots of crosslinks identified in **b** M_i_, **c** M_i_:heparin and **d** M_s_ samples. Colors indicate the frequency of contacts. Dashed boxes highlight N-terminal N1/N2 contacts to P1 and P2/RD. **e** Model of tau conformational changes along the pathway of seed formation show changes in contacts from acidic N-terminal N1/N2 (orange) to P1/P2 (grey) and RD (red, green, blue and magenta).

### Implications for tau aggregation

The structural polymorphisms of tau fibrils are each linked to distinct tauopathies. Thus reagents, such as antibodies or small molecules, specific for different fibrillar structures could be used to accurately diagnose late-stage diseases. Our prior work indicated that small soluble species, including monomer, isolated from either recombinant or tauopathy tissue sources can drive aggregation of inert tau and mediate self-assembly [4]. Structural analysis of recombinant seeds and AD-derived seeds suggested some conserved contacts in their overall topology [4]. More recently, work from Sharma et al [15] revealed that small seeds isolated from disease encode conformational plasticity to maintain a subset of structural polymorphs. It remains unclear how tau seeds form in disease to drive formation of structural polymorphs but it is likely that ligand binding or specific post-translational modifications may promote seed formation to initiate disease. While heparin-induced seeds are likely not physiological, other polyanions such as RNA, may be important ligands that initiate this process [23, 24]. Our data support that heparin binding to the repeat domain of tau displaces specific long-range contacts to the acidic N-terminal N1/N2 domains. Subsequent chromatographic repurification of tau:heparin complexes strips the heparin away allowing the recovery of N-term N1/N2 to P1 contacts but the P2 and RD surfaces remain unmasked which could expose putative amyloid motifs thus promote self-assembly. This model may explain the role of cofactors or PTMs in unmasking specific surfaces for subsequent assembly. It also reveals a role of N-terminal N1/N2 contacts to the proline-rich domain to stabilize monomeric forms of seeds. Additionally, for many years post-translational modifications, such as phosphorylation, were proposed to drive this process [25, 26] but direct evidence for a role of phosphorylation on tau remains unclear. CryoEM structures of tau fibrils isolated from tauopathies suggested the role of acetylation and ubiquitination in the formation of corticobasal degeneration (CBD) and Alzheimer’s disease (AD) structural polymorphs [27, 28] but whether this initiates the process or accumulates once these structures are formed remains an open question. Moreover, disease-derived cryoEM structures of tau have revealed the potential role of yet unknown ligands highlighted by unexplained density in the fibril structures [27, 29]. How seeds relate to structural polymorphs of tau remains elusive but specific exposure of amyloid motifs or ordering of interactions between motifs is likely a central process to mediating tau aggregation. A deeper understanding for how pathogenic seeds are formed and how they drive the formation of distinct structural polymorphs will be required to understand the mechanism of tauopathies and reveal new insight into their treatment.

## Materials and Methods

### Tau purification and SEC of tau species

Wild-type 2N4R tau was purified from *E. Coli* BL21 (DE3) transformed with PET28b plasmid using the same protocol as previously described [3]. The production of tau seeds was carried out by incubating 16uM wild-type tau with 1:1 molar ratio of heparin (AMSbio) for 1 hour at 37°C in 30 mM MOPS pH7.4, 50 mM KCl, 5mM MgCl_2_ and 1 mM with DTT (MOPS buffer). The tau:heparin reactions were injected immediately onto a Superdex 200 Increase 10/300 GL (GE) after incubation in 1X PBS yielding a peak that eluted around 1.5mls earlier than wild-type tau. The seeding activity was confirmed using tau FRET biosensor cells (see below). The bump/dip variation around the bed column volume was normalized to the same buffer. To quantify the remaining heparin in eluted M_s_, heparin Fluorescein (Creative PEGWorks, HP-201 18kDa) alone and in complex with 1:1 molar ratio of 16uM tau were both injected on SEC. The fluorescence intensity from the M_s_ peak was measured by infinite M1000. The standard calibration curve was prepared using the same heparin-FITC in 1X PBS.

### Tau seeding

Samples from SEC elutions were assayed for their seeding activity in HEK293T tau biosensor cells and compared to the same amount of heparin-induced recombinant tau fibril. For all experiments, cells were plated in 96-well plates at 20,000 cells per well in 100 µL of media. 24 hours later, the cells were treated with 30 µL sample:lipofectamine complex. Prior to cell treatment, the recombinant tau fibrils were sonicated for 30 seconds at an amplitude of 65 on a Q700 Sonicator (QSonica). 48 hours after treatment with tau, the cells were harvested by 0.05% trypsin digestion and then fixed in PBS with 2% paraformaldehyde. A BD LSRFortessa was used to perform FRET flow cytometry. To measure mCerulean and FRET signal, cells were excited with the 405 nm laser and fluorescence was captured with a 405/50 nm and 525/50 nm filter, respectively. To measure mClover signal, cells were excited with a 488 laser and fluorescence was captured with a 525/50 nm filter. To quantify FRET, we used a gating strategy where mCerulean bleed-through into the mClover and FRET channels was compensated using FlowJo analysis software. As described previously [3]. FRET signal is defined as the percentage of FRET-positive cells in all analyses. For each experiment, 10,000 cells per replicate were analyzed and each condition was analyzed in triplicate. Data analysis was performed using FlowJo v10 software (Treestar).

### ThT aggregation assay

Wild-type 2N4R was diluted to 17.6 µM in MOPS buffer with 25 μM β-mercaptoethanol and boiled at 100°C for 5 min. A further two-fold dilution in PBS was followed and a final concentration of 25 µM ThT was added in dark. For a 60 µL reaction system, 30 µL tau protein was mixed with equal volume of a mixture consisting of either buffer, seeding monomer at 33nM or 66nM), tau or any combination of them [3]. All experiments were performed in triplicate. ThT kinetic scans were run every 10 min on a Tecan Spark plate reader at 446 nm Ex (5 nm bandwidth), 482 nm Em (5 nm bandwidth) with agitation for 5s prior to acquisition.

### Transmission Electron Microscopy

An aliquot of 5 μL sample was loaded onto a glow-discharged Formvar-coated 300-mesh copper grids for 30 s and was blotted by filter paper followed by washing the grid with 5 μL ddH_2_O. After another 30 seconds, 2% uranyl acetate was loaded on the grids and blotted again. The grid was dried for 1min and loaded into a FEI Tecnai G2 Spirit Biotwin TEM. All images were captured using a Gatan 2Kx2K multiport readout post column CCD at the UT Southwestern EM Core Facility.

### Dynamic Light Scattering aggregation assay

Reactions were prepared using the same experimental conditions as the above ThT fluorescence aggregation experiments. Briefly, wild-type 2N4R was diluted to 17.6 µM in MOPS buffer with 25 uM β-mercaptoethanol and boiled at 100°C for 5 min. A further two-fold dilution in PBS was followed. For a 60 µL reaction volume, 30 µL tau protein was mixed with equal volume of a mixture consisting of either buffer or seeding monomer (with 66nM or 132nM). All protein samples were filtered through a 0.22μm PES sterile filter and loaded in triplicate onto a 384 well clear flat-bottom plate. The plate was loaded into a Wyatt DynaPro Plate Reader III and set to run continuously at room temperature at a scanning rate of 1 scan per 15 minutes, with 1 scan composed of 10 acquisitions for 18 hours. The data were analyzed using the Wyatt Dynamics software version 7.8.2.18. Light scattering results were filtered by Sum of Squares (SOS) <20 to eliminate statistical outlier acquisitions within each scan. Data were reported as average R_h_ over the time course.

### SEC-MALS

2mg/ml of M_i_ alone and in complex with 1:1 molar ratio of heparin (AMSbio) were filtered through a 0.22 μm PES filter before 100 μL each was applied to a Superdex 200 Increase 10/300 column equilibrated in 1xPBS with 1mM TCEP. The column was in line with a Shimadzu UV detector, a Wyatt TREOS II light-scattering detector, and a Wyatt Optilab tREX differential-refractive-index detector. The flow rate was 0.5 mL/min. The data were analyzed with Wyatt’s ASTRA software version 7.1.0.29. SEDFIT was used to calculate the dn/dc of the protein.

### Mass photometry

M_i_ and M_s_ data were acquired in Refeyn OneMP mass photometer. Prior to the measurement, 50uM tau M_i_ was diluted 900X times to 55.5nM and 200nM M_s_ was diluted 30X times to 6.6nM in 1X PBS. 15 µL of each diluted tau sample was injected into the flow-chamber and movies of either 60 or 90 s duration were recorded after autofocus stabilization. Data was processed by Refeyn team through Gaussian fitting to provide the peak mass, the sigma (standard deviation) of the Gaussian, and the number of particles under that Gaussian (and as a % of all counts in the graph). Percentages are with respect to all the particles contained in the graph. Overlapping Gaussian curves will over-count the number of particles in a given population.

### XL-MS of different tau samples

We have developed standardized protocols for crosslinking and data analysis of samples. For DMTMM reactions, protein samples were crosslinked at 0.3mg/ml in 100 µL total volume with a final 12mg/ml DMTMM for 15 minutes at 37°C while shaking at 750rpm. For ADH/DMTMM reactions, protein samples were crosslinked at 0.3mg/ml in 100 µL total volume with a final 8.3mg/ml ADH (d_0_/d_8_, Creative Molecules) and 12mg/ml DMTMM (Sigma-Aldrich) for 15 minutes at 37°C while shaking at 750 rpm. The reactions were quenched with 200mM Ammonium Bicarbonate (AB) for 30 minutes. Samples were resolved on SDS-PAGE gels (NUPAGE™, 4 to 12%, Bis-tris, 1.5mm or home-made SDS-Gel) and bands corresponding to tau monomer were gel-extracted following standard protocols [3]. Samples were flash frozen in liquid nitrogen, lyophilized and resuspended in 8M urea followed by 2.5mM TCEP reduction and 5mM Iodoacetamide alkylation in dark with each 30 minutes. Samples were then diluted to 1M urea by 50mM AB and digested by 1:50 (m/m) trypsin (Promega) overnight shaking at 600rpm. 2% (v/v) formic acid was added to acidify the reaction system and further purified by reverse-phase Sep-Pak tC18 cartridges (Waters) and size exclusion peptide chromatography (SEPC). Fraction collected from SEPC was lyophilized. The dried samples were resuspended in water/acetonitrile/formic acid (95:5:0.1, v/v/v) to a final concentration of approximately 0.5 µg/µL. 2 µL of each was injected into Eksigent 1D-NanoLC-Ultra HPLC system coupled to a Thermo Orbitrap Fusion Tribrid system at the UTSW Proteomics core.

The analysis of the mass spectrum data was done by in-house version of xQuest [30]. Each Thermo.raw data was first converted to open.mzXML format using mscovert (proteowizard.sourceforge.net). Search parameters were set differently based on the crosslink reagent as followed. For DMTMM zero-length crosslink search: maximum number of missed cleavages = 2, peptide length = 5-50 residues, fixed modifications = carbamidomethyl-Cys (mass shift = 57.02146 Da), mass shift of crosslinker = -18.010595 Da, no monolink mass specified, MS1 tolerance = 15 ppm, and MS2 tolerance = 0.2 Da for common ions and 0.3 Da for crosslink ions; search in enumeration mode. For ADH, maximum number of missed cleavages (excluding the crosslinking site) = 2, peptide length = 5–50 residues, fixed modifications = carbamidomethyl-Cys (mass shift = 57.021460 Da), mass shift of the light crosslinker = 138.09055 Da, mass shift of monolinks = 156.10111 Da, MS1 tolerance = 15 ppm, MS2 tolerance = 0.2 Da for common ions and 0.3 Da for crosslink ions, search in ion-tag mode. FDRs were estimated by xprophet [31] to be 0% -0.17%. For ADH For each experiment, five replicate data sets were compared and the frequency of contacts were calculated. The pairs position and unique nseen numbers (frequencies) were visualized using custom gunplot script.

## Acknowledgements

This work was supported by grants to L.A.J from the Marie Effie Cain Endowed Scholarship, a Chan Zuckerberg Initiative Collaborative Science Award (2018-191983) and a Bright Focus Foundation grant (A2019060). We appreciate the help of the Molecular Biophysics Resource core, Structural Biology Laboratory, Cryo-Electron Microscopy Facility and Proteomics Core Facility at the University of Texas Southwestern Medical Center. We also thank members of the Joachimiak lab for reading and providing critical comments on the manuscript.

## Author Contributions

Z.H. and L.A.J. conceived and designed the overall study. Z.H. purified all the recombinant proteins, developed method to isolate M_s_, acquired and analyzed DLS, SEC-MALS and mass photometry data. D.C. acquired and analyzed all crosslinking experiments. B.D.R collected TEM images of samples. Z.H., D.C. and L.A.J. wrote the manuscript, and all authors contributed to its improvement.

## Data availability

The data sets generated during and/or analyzed during the current study are available from the corresponding authors on reasonable request.

## SUPPLEMENTARY INFORMATION

## SUPPLEMENTARY FIGURES AND LEGENDS

**Supplementary Figure 1.**
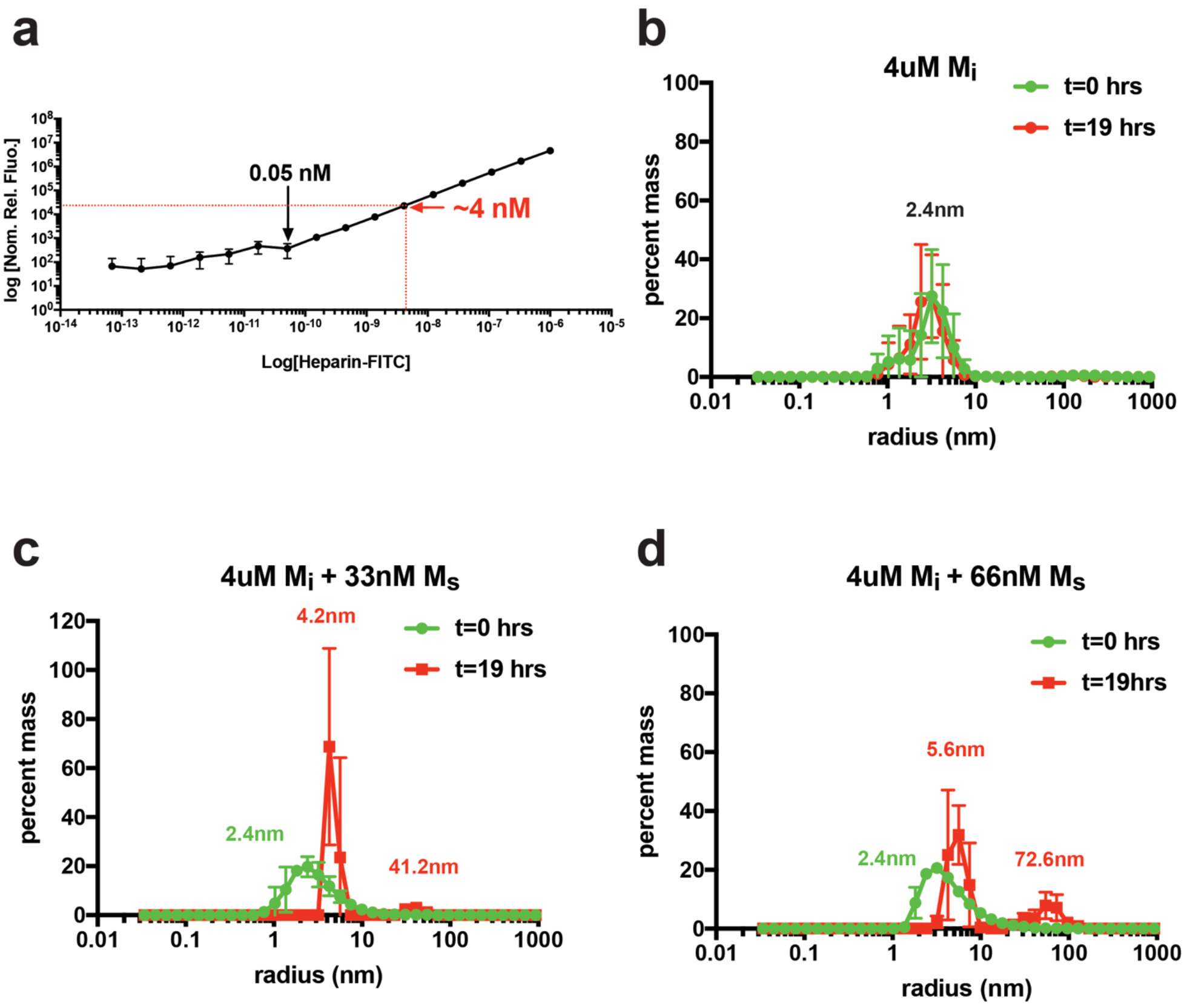
Assembly properties of a tau seed. **a** Calibration curve of FITC fluorescence using excitation 495nm and emission 520nm. Signal detected in the M_i_:heparin-FITC reaction from the M_s_ fraction corresponds to 4 nM FITC. **b** Histogram of size distributions of 4uM M_i_ at times 0 hour (green) and 19 hour (red). Median size is indicated above the distribution. **c** Histogram of size distributions of 4uM M_i_ with 33 nM M_s_ at times 0 hour (green) and 19 hour (red). **d** Histogram of size distributions of 4uM M_i_ with 66 nM M_s_ at times 0 hour (green) and 19 hour (red). Median species size is indicated above the distribution and is colored green for 0 hour and red for 19 hour. Each DLS experiment was performed in triplicate and the data is shown as averages with standard deviation.

**Supplementary Figure 2.**
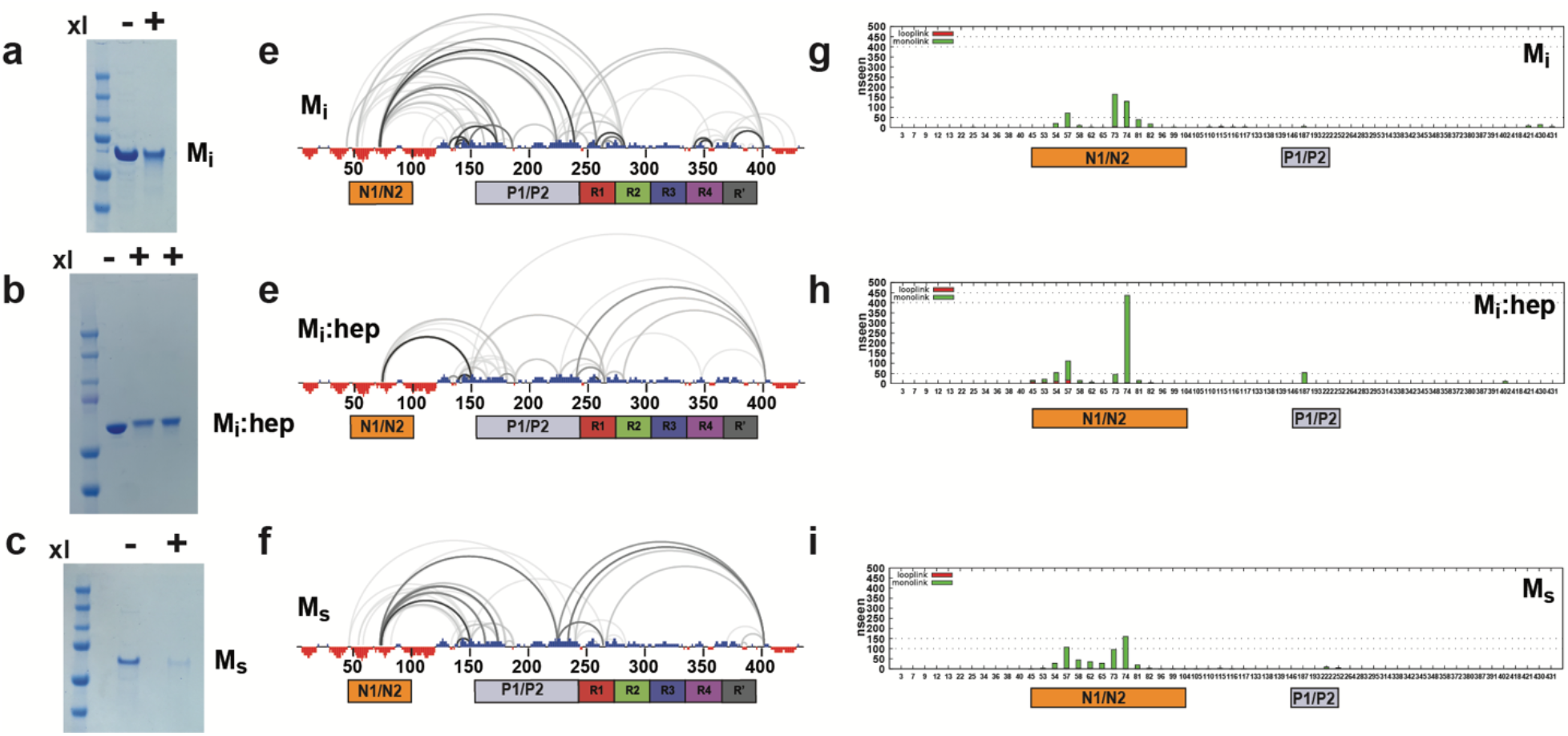
**a-c** M_i_, M_i_:heparin and M_s_ samples were reacted with DMTMM/ADH for 15 minutes and the non-crosslinked and crosslinked reactions were resolved on SDS-PAGE. Linear plots illustrating the distribution of zero-length contacts in **d** M_i_, **e** M_i_:heparin and **f** M_s_ across 5 replicate samples. The crosslinks are shown as semi-circles and are colored according to frequency in the experiments. Cartoon schematic for 2N4R tau highlighting the N1/N2, P1/P2 and repeat domains colored as in Fig 1a. **g-i** Frequencies of monolinks (green) and looplinks (red) derived from the M_i_, M_i_:heparin and M_s_ crosslinking reactions. Modifications are derived from ADH reactions and are shown as a bar plot using the mean frequency of monolinks and looplinks across 5 replicates. ADH monolinks and looplinks are predominantly observed in the acidic N-terminal N1/N2 (orange) and their relative frequency can be interpreted as solvent accessibility. Only minor changes are observed in the P1/P2 region (grey).

## References

1. Iqbal, K., et al., Tau in Alzheimer disease and related tauopathies. Curr Alzheimer Res, 2010. 7(8): p. 656–64.

2. Levine, Z.A., et al., Regulation and aggregation of intrinsically disordered peptides. Proc Natl Acad Sci U S A, 2015. 112(9): p. 2758–63.

3. Chen, D., et al., Tau local structure shields an amyloid-forming motif and controls aggregation propensity. Nat Commun, 2019. 10(1): p. 2493.

4. Mirbaha, H., et al., Inert and seed-competent tau monomers suggest structural origins of aggregation. Elife, 2018. 7.

5. Mocanu, M.M., et al., The potential for beta-structure in the repeat domain of tau protein determines aggregation, synaptic decay, neuronal loss, and coassembly with endogenous Tau in inducible mouse models of tauopathy. J Neurosci, 2008. 28(3): p. 737–48.

6. Kar, S., et al., Repeat motifs of tau bind to the insides of microtubules in the absence of taxol. EMBO J, 2003. 22(1): p. 70–7.

7. Wolfe, M.S., Tau mutations in neurodegenerative diseases. J Biol Chem, 2009. 284(10): p. 6021–5.

8. Agerschou, E.D., et al., beta-Turn exchanges in the alpha-synuclein segment 44-TKEG-47 reveal high sequence fidelity requirements of amyloid fibril elongation. Biophys Chem, 2021. 269: p. 106519.

9. Ramachandran, G. and J.B. Udgaonkar, Understanding the kinetic roles of the inducer heparin and of rod-like protofibrils during amyloid fibril formation by Tau protein. J Biol Chem, 2011. 286(45): p. 38948–59.

10. Meraz-Rios, M.A., et al., Tau oligomers and aggregation in Alzheimer’s disease. J Neurochem, 2010. 112(6): p. 1353–67.

11. Zhu, H.L., et al., Quantitative characterization of heparin binding to Tau protein: implication for inducer-mediated Tau filament formation. J Biol Chem, 2010. 285(6): p. 3592–9.

12. Fichou, Y., et al., Cofactors are essential constituents of stable and seeding-active tau fibrils. Proc Natl Acad Sci U S A, 2018. 115(52): p. 13234–13239.

13. Zhang, W., et al., Heparin-induced tau filaments are polymorphic and differ from those in Alzheimer’s and Pick’s diseases. Elife, 2019. 8.

14. Kundu, B., et al., Nucleation-dependent conformational conversion of the Y145Stop variant of human prion protein: structural clues for prion propagation. Proc Natl Acad Sci U S A, 2003. 100(21): p. 12069–74.

15. Sharma, A.M., et al., Tau monomer encodes strains. Elife, 2018. 7.

16. Nachman, E., et al., Disassembly of Tau fibrils by the human Hsp70 disaggregation machinery generates small seeding-competent species. J Biol Chem, 2020. 295(28): p. 9676–9690.

17. Ohhashi, Y., et al., Molecular basis for diversification of yeast prion strain conformation. Proc Natl Acad Sci U S A, 2018. 115(10): p. 2389–2394.

18. Abskharon, R., et al., Crystal structure of a conformational antibody that binds tau oligomers and inhibits pathological seeding by extracts from donors with Alzheimer’s disease. J Biol Chem, 2020. 295(31): p. 10662–10676.

19. Kaufman, S.K., et al., Characterization of tau prion seeding activity and strains from formaldehyde-fixed tissue. Acta Neuropathol Commun, 2017. 5(1): p. 41.

20. Furman, J.L., et al., Widespread tau seeding activity at early Braak stages. Acta Neuropathol, 2017. 133(1): p. 91–100.

21. Holmes, B.B., et al., Proteopathic tau seeding predicts tauopathy in vivo. Proc Natl Acad Sci U S A, 2014. 111(41): p. E4376–85.

22. Hou, Z., et al., DnaJC7 binds natively folded structural elements in tau to inhibit amyloid formation. BioRxiv, 2020.

23. Kampers, T., et al., RNA stimulates aggregation of microtubule-associated protein tau into Alzheimer-like paired helical filaments. FEBS Lett, 1996. 399(3): p. 344–9.

24. Dinkel, P.D., et al., RNA Binds to Tau Fibrils and Sustains Template-Assisted Growth. Biochemistry, 2015. 54(30): p. 4731–40.

25. Park, S., et al., Degradation or aggregation: the ramifications of post-translational modifications on tau. BMB Rep, 2018. 51(6): p. 265–273.

26. Haj-Yahya, M. and H.A. Lashuel, Protein Semisynthesis Provides Access to Tau Disease-Associated Post-translational Modifications (PTMs) and Paves the Way to Deciphering the Tau PTM Code in Health and Diseased States. J Am Chem Soc, 2018. 140(21): p. 6611–6621.

27. Arakhamia, T., et al., Posttranslational Modifications Mediate the Structural Diversity of Tauopathy Strains. Cell, 2020. 180(4): p. 633–644 e12.

28. Fitzpatrick, A.W.P., et al., Cryo-EM structures of tau filaments from Alzheimer’s disease. Nature, 2017. 547(7662): p. 185–190.

29. Falcon, B., et al., Novel tau filament fold in chronic traumatic encephalopathy encloses hydrophobic molecules. Nature, 2019. 568(7752): p. 420-423.

30. Rinner, O., et al., Identification of cross-linked peptides from large sequence databases. Nat Methods, 2008. 5(4): p. 315–8.

31. Walzthoeni, T., et al., False discovery rate estimation for cross-linked peptides identified by mass spectrometry. Nat Methods, 2012. 9(9): p. 901–3.

